# The protein-bound uremic solute 3-carboxy-4-methyl-5-propyl-2-furanpropionate (CMPF) increases erythrocyte osmotic fragility through interaction with the mechanosensitive channel Piezo1

**DOI:** 10.1101/2023.05.26.542480

**Authors:** Beatriz Akemi Kondo Van Spitzenbergen, Gabriela Bohnen, Erika Sousa Dias, Júlia B. Monte Alegre, Gabriela Ferreira Dias, Nadja Grobe, Andrea Novais Moreno-Amaral, Peter Kotanko

## Abstract

Jedi1, a chemical activator of the mechanosensitive cation channel Piezo1, shares structural similarities with the uremic solute 3-carboxy-4-methyl-5-propyl-2-furanpropionate (CMPF). We explored the hypothesis that CMPF at a concentration seen in uremia activates Piezo1 located on red blood cells (RBC). We incubated RBC from five healthy individuals with either Jedi1 or CMPF (both 87 μM), with or without the Piezo1 inhibitor GsMTx-4 (2 μM), and quantified the cells’ osmotic fragility. Our results indicate that compared to controls (i.e., RBC incubated with buffer), both Jedi1 and CMPF increase the osmotic fragility of RBC (i.e., reduce their resistance to osmotic stress). Effects of Jedi1 and CMPF were reversed to the control level by GsMTx-4. These results indicate a role of Piezo1 in augmenting RBC osmotic fragility and modulation of this effect by Jedi1 and CMPF. Our findings open the possibility that CMPF may act as an endogenous chemical activator of Piezo1.

## Introduction

Anemia is common in individuals with chronic kidney disease (CKD). Insufficient renal production of erythropoietin is considered the main cause of renal anemia. Several additional factors contribute to renal anemia, such as functional iron deficiency due to chronic inflammation and shortened life span of red blood cells (RBC) due to eryptosis. In CKD patients treated with hemodialysis, several groups reported a shortened RBC life span (RBCLS) from 100 to 120 days observed in healthy individuals to 50 to 100 days^1, 2, 3^. While the etiology of shortened RBCLS is ill-defined, the “uremic milieu” is considered a key factor^4, 5^. Eryptosis can be induced by the uremic retention solutes methylglyoxal and indoxyl sulfate^6, 7^, and osmotic^8^ and oxidative stress^9^. Eryptotic RBC show increased levels of intracellular Ca^2+^ ^8^, reactive oxygen species (ROS)^10^, and decreased ATP levels^11^. Eryptotic events are triggered by the influx of Ca^2+^ through selective cation channels^12, 13^, which stimulates the translocation of phosphatidylserine to the RBC outer membrane leaflet^11^. Phosphatidylserine on the RBC surface facilitates the recognition, phagocytosis, and degradation of RBC by macrophages^11^ and pro-inflammatory monocytes^14^.

Recently, we have hypothesized that Ca^2+^ influx through the mechanosensitive cation channel Piezo1 located in the RBC plasma membrane may be involved in the genesis of eryptosis^15^. Piezo1 plays a critical role in RBC volume regulation^16^, whereby shear stress in the microcirculation activates Piezo1-dependent Ca^2+^ influx. The ensuing increase of intracellular Ca^2+^ subsequently activates calcium-activated potassium KCa3.1 Gardos channels^17^, leading to hyperpolarization and loss of K^+^, Cl^−^, and water, resulting in RBC shrinkage. In addition, forming the calcium-calmodulin complex destabilizes the actin-adducin-Band 4.1 complex, which makes the cross-linked spectrin network more flexible^18^. This process is physiologically important as the RBC traverses narrow anatomical structures such as capillaries and the slits that are formed in the spleen by the spaces between the endothelial cells lining the organ’s blood vessels. After the passage, both Piezo1 and the Gardos-channels are inactivated and return to their respective closed states. Na^+^/K^+^ATPase and Ca^2+^ATPase pumps re-establish pre-passage intracellular ion concentrations, allowing RBC to return to its original size^18^. While mechanical stress is the only known physiological activator of Piezo1, four synthetic molecules, Yoda1, Yoda2, Jedi1, and Jedi2 (the latter two also abbreviated as Jedi1/2), have been identified as chemical Piezo1 activators. These molecules increase the open probability of Piezo1 in the absence of mechanical stimulation^19, 20^. While Yoda1 and Yoda2 do not share structural similarities with Jedi1/2, all four molecules activate Piezo1 in a dose-dependent fashion^20^.

Functional studies suggest that the molecules act by different mechanisms: while Jedi1/2 activate Piezo1 through binding to its extracellular domain in its upstream blade at residues L15-16 and L19-20, Yoda1 and Yoda2 act through binding to the downstream beam. Research into mechanosensitive ion channels has benefitted from the discovery of GsMTx-4, a gating modifier peptide from spider venom^21, 22^, that selectively inhibits cation-permeable mechanosensitive channels, including Piezo1^23, 24^. GsMTx-4 acts from the extracellular side and occupies a small fraction of the surface area in the unstressed plasma membrane adjacent to Piezo1. When applied tension reduces lateral pressure in the lipids, GsMTx-4 penetrates deeper and acts as an ‘‘area reservoir’’ leading to partial relaxation of the outer membrane layer and thus decreasing the effective magnitude of mechanical stimulus^25, 26^.

Notably, Jedi1 (2-methyl-5-phenylfuran-3-carboxylic acid) and Jedi2 (2-methyl-5-(2-thienyl)-3-furancarboxylic acid) share the Piezo1-activating moiety, 3-carboxylic acid methylfuran^20^. The functional relevance of this moiety has been demonstrated in experiments where *N*-butylaniline, a molecule lacking this structure, did not activate Piezo1^20^. Kotanko *et al*. (2022) recently observed that the active moiety of Jedi1/2 shows structural similarity with 3-carboxy-4-methyl-5-propyl-2-furanpropanoic acid (CMPF) (**Figure 1**), a uremic retention solute^15^. In CKD patients treated with hemodialysis, CMPF levels are increased 5- to 15-fold compared to healthy individuals^27^. Given this structural resemblance, Kotanko *et al*. (2022) hypothesized that increased levels of CMPF lengthen the Piezo1 activation in a Jedi1/2-like fashion. The authors further hypothesized that this CMPF-induced activation might trigger a cascade of events that eventually results in eryptosis and shortened RBCLS ^15^.

**Figure 1.**
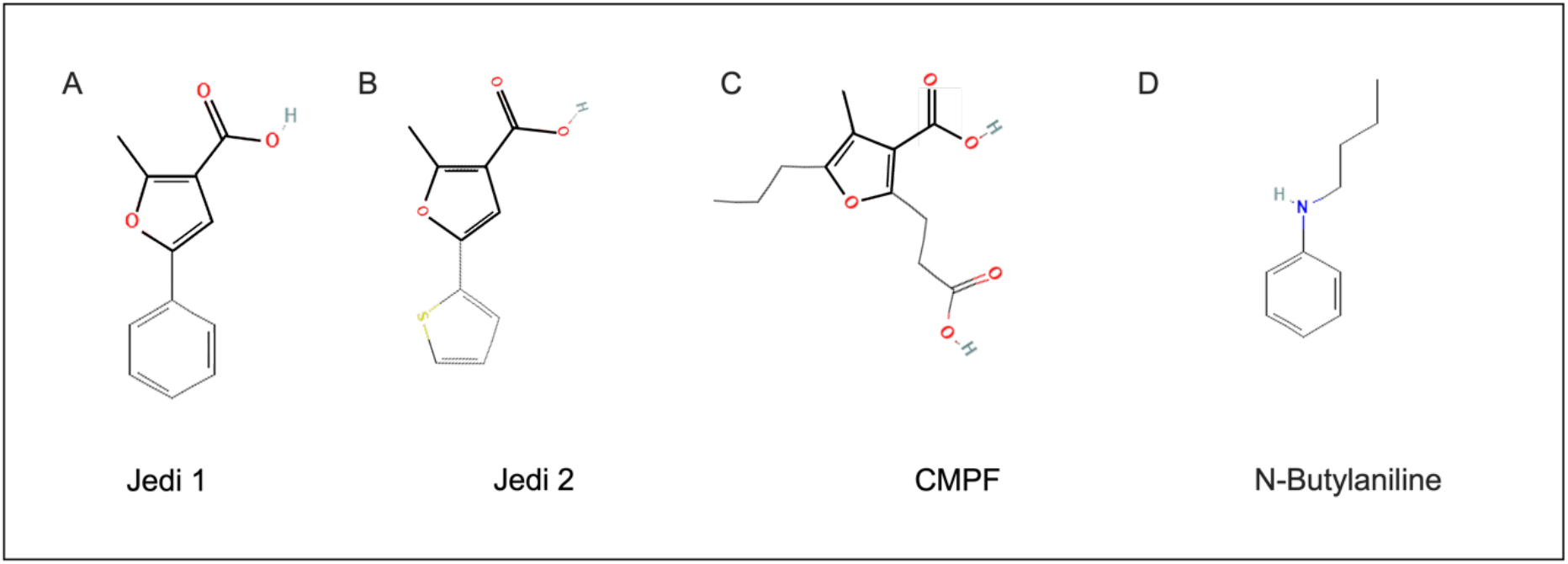
Two-dimensional structures of compounds. (A) Jedi1 (2-Methyl-5-phenylfuran-3-carboxylic acid), a chemical activator of Piezo1. (B) Jedi2 (2-Methyl-5-(thien-2-yl)-3-furoic acid), a chemical activator of Piezo1. (C) CMPF (3-Carboxy-4-methyl-5-propyl-2-furanpropionic acid), a protein-bound uremic retention solute derived from furan fatty acid metabolism. CMPF has been hypothesized to activate Piezo1 in a Jedi1/2-like fashion due to their shared 3-carboxylic acid methylfuran moiety, the putative Piezo1 activating domain; this structure is highlighted in A, B, and C^15^. (D) N-Butylaniline. This molecule lacks the 3-carboxylic acid methylfuran moiety and did not activate Piezo1 in experiments conducted by Wang et al. (2018) ^20^. Source: PubChem.

We explored the hypothesis that CMPF at a concentration seen in uremia activates Piezo1 located on RBC. To that end, we incubated RBC from healthy subjects with Jedi1 or CMPF in the presence or absence of GsMTx-4 and measured their osmotic resistance.

## Results

### Study population

RBC were obtained from five healthy subjects (two males, three females). Their mean±SD age was 28.2±6.57 years. Baseline characteristics are presented in **Table 1**.

**Table 1.** Demographical and laboratory characteristics of the study population.

| Parameters | Healthy subjects (n = 5) |
| --- | --- |
| Age (years) | 28.20±6.57 |
| Sex | 2 males; 3 females |
| Caucasians | 5 |
| Body mass index (kg/m<sup>2</sup>) | 22.85±3.07 |
| Hemoglobin (g/dL) | 14.20±1.90 |
| Creatinine (mg/dL) | 0.94±0.21 |
| Urea (mg/dL) | 27.20±4.71 |
| Albumin (g/dL) | 4.24±0.33 |

Data expressed as mean±SD.

### Osmotic fragility test

The five panels in **Figure 2** show RBC osmotic fragility curves (OFC) from a single donor under different experimental conditions. **Figure 2A** shows the OFC from the negative control (RBC incubated in buffer). The osmotic fragility index (OFI) was 4.77±0.03 g/L. In the presence of Jedi1 (87 μM), a right-shift of the OFC was observed (**Figure 2B**; OFI 5.09±0.02 g/L), indicating an increased osmotic fragility. This effect of Jedi1 was not observed in the presence of Piezo1 inhibitor GsMTx-4 (2 μM) (**Figure 2C**; OFI 4.35±0.06 g/L). Incubation of RBC with CMPF (87 μM) resulted in a marked right-shift of the OFC (**Figure 2D**; OFI 5.83±0.04 g/L). Incubation with CMPF in the presence of GsMTx-4 showed an OFI comparable to negative control (**Figure 2E**; OFI 4.36±0.02 g/L).

**Figure 2.**
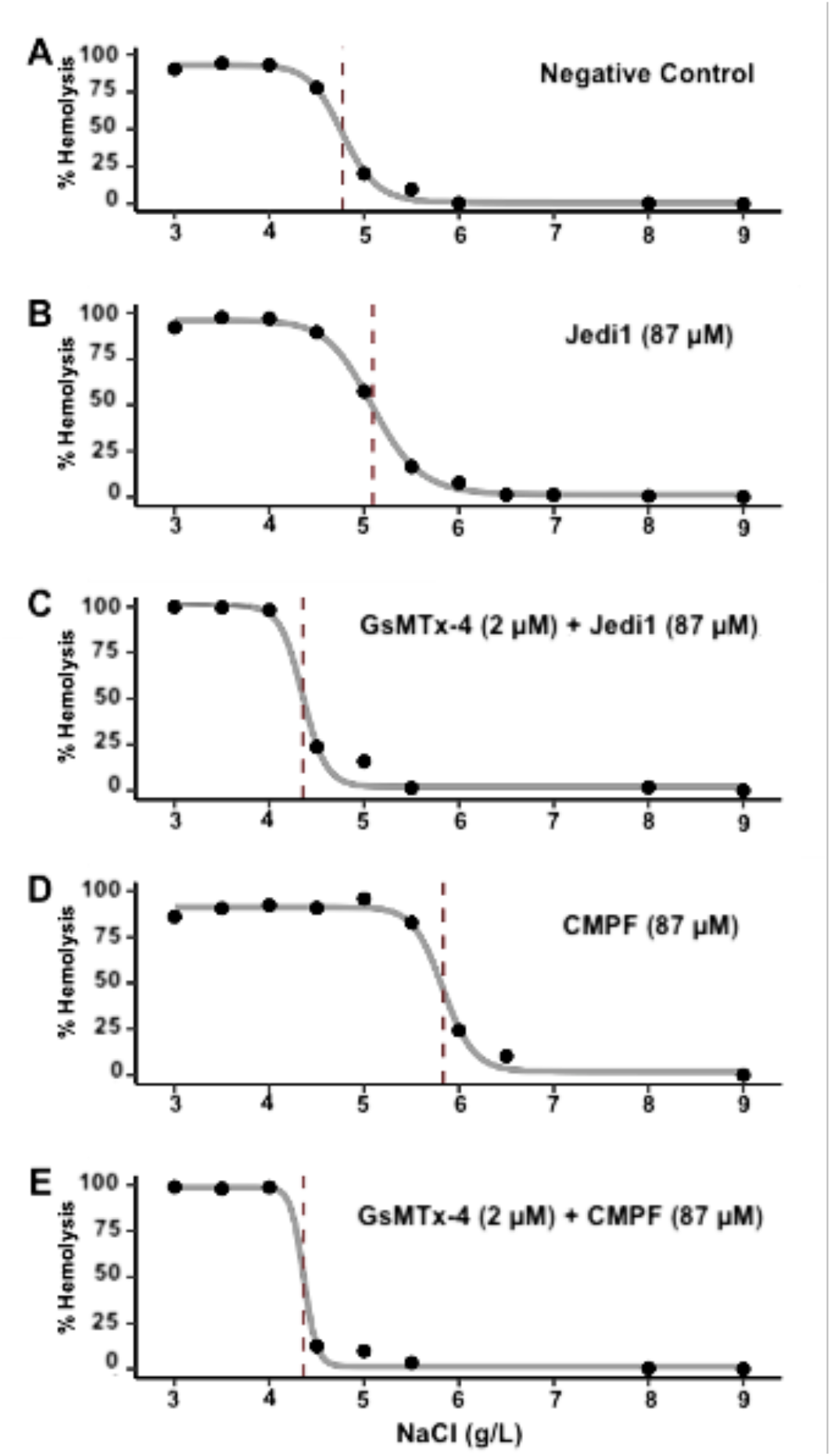
Representative red blood cell (RBC) osmotic fragility curves. The dashed lines indicate the osmotic fragility index (OFI). (A) Negative control consisted of RBC incubated in incubation buffer (phosphate buffered saline, 4% human serum albumin, and 0.12% DMSO). (B) RBC incubated with Jedi1 (87 μM); (C) RBC incubated with GsMTx-4 (2 μM) + Jedi1 (87 μM); (D) RBC incubated with CMPF (87 μM); (E) RBC incubated with GsMTx-4 (2 μM) + CMPF (87 μM). Jedi1, CMPF, and GsMTx-4 were dissolved in the incubation buffer.

The results from all five donors are shown in **Figure 3** and **Table 2**. Incubation with Jedi1 resulted in a significantly higher mean OFI than that in control (5.02±0.28 g/L vs. 4.57±0.31 g/L, respectively). The same effect was not observed when cells were pre-treated with GsMTx-4, as the mean OFI was not different than that of the control (4.56±0.29 g/L vs. 4.57±0.31 g/L, respectively). When RBCs were incubated with CMPF, the OFI values of all donors increased significantly (5.57±0.39 g/L). Interestingly, the use of a mechanosensitive channel inhibitor prior to uremic toxin treatment prevented the CMPF-induced increase of OFI compared with the control (4.47±0.13 g/L vs. 4.57±0.31 g/L, respectively).

**Table 2.**
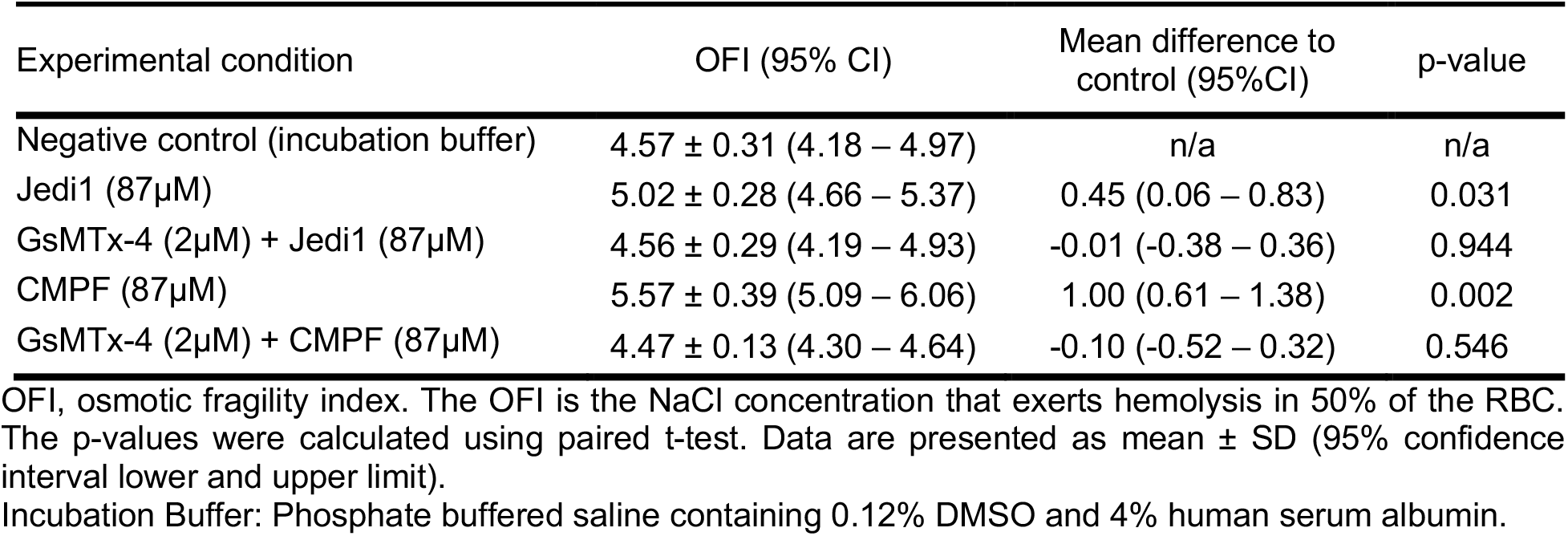
Osmotic fragility index of RBC under different experimental conditions.

| Experimental condition | OFI (95% CI) | Mean difference to control (95%CI) | p-value |
| --- | --- | --- | --- |
| Negative control (incubation buffer) | 4.57 ± 0.31 (4.18 – 4.97) | n/a | n/a |
| Jedi1 (87µM) | 5.02 ± 0.28 (4.66 – 5.37) | 0.45 (0.06 – 0.83) | 0.031 |
| GsMTx-4 (2µM) + Jedi1 (87µM) | 4.56 ± 0.29 (4.19 – 4.93) | -0.01 (-0.38 – 0.36) | 0.944 |
| CMPF (87µM) | 5.57 ± 0.39 (5.09 – 6.06) | 1.00 (0.61 – 1.38) | 0.002 |
| GsMTx-4 (2µM) + CMPF (87µM) | 4.47 ± 0.13 (4.30 – 4.64) | -0.10 (-0.52 – 0.32) | 0.546 |

OFI, osmotic fragility index. The OFI is the NaCl concentration that exerts hemolysis in 50% of the RBC. The p-values were calculated using paired t-test. Data are presented as mean ± SD (95% confidence interval lower and upper limit).
Incubation Buffer: Phosphate buffered saline containing 0.12% DMSO and 4% human serum albumin.

**Figure 3.**
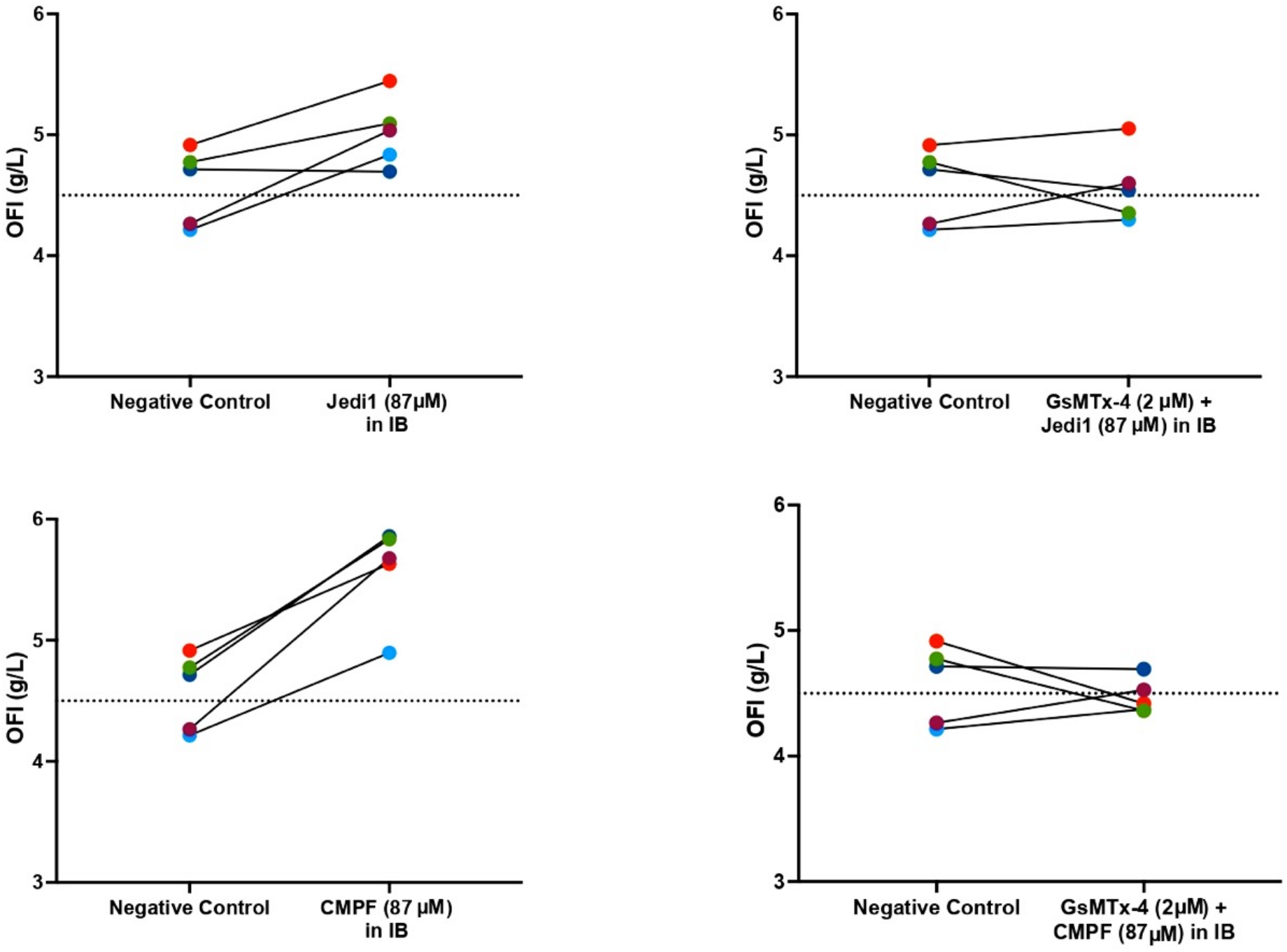
Osmotic Fragility Index (OFI) expressed as the concentration of NaCl (in g/L) that exerted 50% hemolysis. The individual donors are color-coded, the experimental conditions are indicated on the x-axis. Solid lines connect OFI results obtained under control and incubation conditions, respectively. The horizontal dotted lines indicate the mean OFI in controls. Legend: IB - Incubation buffer. The incubation buffer comprised phosphate buffered saline, 4% human serum albumin, and 0.12% DMSO. and served as negative control.

## Discussion

Piezo1 is a phylogenetically old and highly conserved mechanosensitive cation channel located on several cell types, including mammalian RBC^28, 29^. Physiologically, Piezo1 is activated by mechanical stimulation. While four exogenous, synthetic chemical activators of Piezo1 have been described (Jedi1, Jedi2, Yoda1, Yoda2), we are not aware of endogenous chemical activators. Because CMPF, a furan fatty acid metabolite that accumulates in CKD patients, shares structural similarities with Jedi1/2, it has been hypothesized by Kotanko *et al*. (2022) that CMPF, at concentrations seen in patients with kidney failure, may act as an endogenous chemical activator of Piezo1 located on RBC surface and instigate eryptosis^15^. Our research presented herein provides the first evidence supporting this hypothesis.

In our experiments, we incubated RBC from healthy donors with either Jedi1 or CMPF and in the presence or absence of the Piezo1 inhibitor GsMTx-4. RBC were then exposed to hypotonic stress. Cell swelling induced by hypotonic solution is an experimental model of *in vitro* Piezo1 activation^18, 30^. Increased plasma membrane tension induced by cell swelling most likely results in mechanical activation^31^.The readouts of interest were the osmotic fragility curve (OFC) and the through curve-fitting derived osmotic fragility index (OFI), an established indicator of RBC osmotic fragility. OFC has been used to assess Piezo1 activation in isolated RBC from mice^16^ and humans^30^. Our experiments revealed that incubation of isolated RBC with either Jedi1 or CMPF resulted in a right-shift of the OFC and a significant increase in OFI. The effects of Jedi1 and CMPF were mitigated by the Piezo1 antagonist GsMTx-4.

RBC incubated for 30 minutes with Jedi1 at concentrations of 100 μM, 300 μM, and 1 mM exhibited a slightly augmented hemolysis when subsequently exposed to hypotonic stress^30^. These findings are corroborated by our results, in which osmotically stimulated RBC showed enhanced hemolysis when treated with 87 μM Jedi1. Surface resonance binding assays revealed that Jedi1 acts through the extracellular regions of the peripheral blade of Piezo1^32, 20^. Experiments conducted by Wang et al (2018) highlight the importance of the 3-carboxylic acid methylfuran domain for the activation of Piezo1. The authors found that N-Butylaniline, a compound lacking this moiety, failed to bind and activate Piezo1^30^. Of note, this 3-carboxylic acid methylfuran domain is shared by Jedi1/2 and CMPF^15^ (Kotanko et al. 2022). Taken together, these results open the possibility that CMPF interacts with Piezo1 in a Jedi1-like manner. However, understanding how exactly CMPF interacts with Piezo1 warrants further investigation.

If CMPF indeed delays Piezo1 inactivation, Ca^2+^ influx would be prolonged. The importance of the extended Ca^2+^ influx has been described in Piezo1 gain-of-function (GOF) mutations that delay channel inactivation. GOF mutations are observed in patients with dehydrated hereditary stomatocytosis, a non-immune congenital hemolytic disorder^33, 34^. In sickle cell disease, the characteristic RBC shape has been attributed to cell dehydration^35^. Of note, RBC sickling can be inhibited by the Piezo1 blocker GsMTx-4^36^. Our study has some limitations apart from the small sample size. Measuring intracellular Ca^2+^ concentration following exposure to CMPF will add to our understanding of the interaction between CMPF and Piezo1. In addition, quantifying phosphatidylserine exposure upon RBC incubation with CMPF will help elucidate the link between CMPF and eryptosis.

In summary, our results indicate the possibility that CMPF activates Piezo1 in RBC from healthy subjects. A detailed understanding of the molecular interaction between CMPF and Piezo1 warrants future studies. By revealing CMPF as a potential endogenous Piezo1 agonist, the present work adds new perspectives to our understanding of this mechanoreceptor. In reference to patients with CKD, understanding the molecular mechanisms of the interaction between CMPF and Piezo1 could instigate novel therapeutic avenues.

## Methods

This study was approved by the ethics committee of Pontifícia Universidade Católica do Paraná (registration number 5.697.460), and all participants gave written informed consent before blood collection.

### Chemicals and Solutions

CMPF and Jedi1 (Sigma-Aldrich, MO, USA) were dissolved in dimethyl sulfoxide (DMSO). A literature review was conducted to determine the concentration of CMPF used for the incubation (data not shown). Each compound was tested at a concentration of 87 μM in 0.12% DMSO. Phosphate buffered saline (PBS) containing 0.12% of DMSO and 4% of human serum albumin (HSA) (Sigma-Aldrich, MO, USA) was used as the negative control and incubation buffer. The mechanosensitive ion channel inhibitor GsMTx-4 (Abcam, MO, USA) was dissolved in PBS and tested at 2 μM.

### Subjects and blood samples

Healthy subjects (n = 5) without a history of renal or inflammatory diseases and who did not receive anti-inflammatory medication or blood transfusion 3 months prior to blood draw were selected. Venous blood was drawn into 3.2% sodium citrate tubes (BD^®^ Vacutainer, BD Biosciences, NJ, USA). For erythrocyte isolation, whole blood was centrifuged (3000 rpm, 15 min, 4°C), the plasma and buffy coat were discarded, and cells were washed twice with cold PBS.

### Osmotic Fragility Test

A volume of 200 μL of isolated RBC was suspended in 800 μL of PBS 4% HSA (20% final hematocrit). This suspension was then incubated with or without 2 μM of GsMTx-4 at room temperature (RT). After 5 minutes, 1.25 μL of CMPF or Jedi1 stocks (both 69.6 mM) or DMSO were added to the suspensions and incubated for 30 minutes at RT. 10 μL of the suspension were transferred to 1 mL of double distilled water, representing total hemolysis, or increasing concentrations of NaCl solutions (in g/L: 3, 3.5, 4, 4.5, 5, 5.5, 6, 6.5, 7, 8, 9) diluted in 1 mL final volume of double distilled water containing CMPF, Jedi1 (both 87 M), or DMSO (0.12%), and incubated for 5 minutes at RT. Each experimental condition was replicated two times.

The content of each saline tube was centrifuged at 1500 × rpm for 10 min, and 250 μL of the supernatant was subjected to hemoglobin optical density (OD) determination at 540 nm (Molecular Devices, CA, USA). The percentage of hemolysis for each NaCl concentration was calculated as follows:

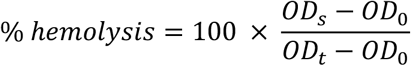

where OD_s_ = OD of the supernatant from RBC subjected to NaCl; OD_0_ = OD of the supernatant obtained from the non-lysed negative control RBC (NaCl 9 g/L); and OD_t_ = OD of supernatant from RBC subjected to total hemolysis^37^. The osmotic fragility index (OFI) was defined as the NaCl concentration that exerted 50% hemolysis. Some authors refer to OFI as C50^16^ or LC50^30^.

### Statistical analysis

OFI and osmotic fragility curves (OFC) were calculated using R version 4.1.1 with packages drc and ggplot2 (R Foundation for Statistical Computing, Vienna, Austria). The OFI distribution was evaluated through the Kolmogorov-Smirnov test. Paired t-test (GraphPad Prism Version 9) was used to compare OFI between control and incubations.

## Acknowledgments

The donation of biological samples from healthy subjects is greatly appreciated by the authors. This research was supported by Renal Research Institute, New York, NY, USA grants. Spitzenbergen, Beatriz Akemi Kondo Van (master student) and Bohnen, Gabriela. (Ph.D. student) are the recipients of research fellowships from Conselho Nacional de Desenvolvimento Científico e Tecnológico (CNPq) and the Coordenação de Aperfeiçoamento de Pessoal de Nível Superior (CAPES).

## Notes

### Competing Interest Statement

The authors have declared no competing interest.

